# ASSEMBLY AND DYNAMICS OF MICROBIAL COMMUNITIES IN GRANULAR, FIXED-BIOFILM AND PLANKTONIC METHANOGENIC MICROBIOMES VALORIZING LONG CHAIN FATTY ACID (LCFA)-RICH WASTEWATER

**DOI:** 10.1101/2021.07.29.454224

**Authors:** Suniti Singh, Johanna M. Rinta-Kanto, Piet N. L. Lens, Marika Kokko, Jukka Rintala, Vincent O’Flaherty, Umer Zeeshan Ijaz, Gavin Collins

## Abstract

Distinct microbial assemblages are engineered in anaerobic digestion (AD) reactors to drive sequential conversions of organics to methane. The spatio-temporal development of three such assemblages (granules, biofilms, planktonic) derived from the same inoculum was studied in replicated bioreactors treating long-chain fatty acids (LCFA)-rich wastewater at 20°C at hydraulic retention times (HRTs) of 12-72 h. We found granular, biofilm and planktonic assemblages differentiated by diversity, structure, and assembly mechanisms; demonstrating a spatial compartmentalisation of the microbiomes from the initial community reservoir. Our analysis linked abundant *Methanosaeta* and *Syntrophaceae*-affiliated taxa (*Syntrophus* and *uncultured*) to their putative, active roles in syntrophic LCFA bioconversion. LCFA loading rates (stearate, palmitate), and HRT, were significant drivers shaping microbial community dynamics and assembly. This study of the archaea and syntrophic bacteria actively valorising LCFAs at short HRTs and 20°C will help uncover the microbiology underpinning anaerobic bioconversions of fats, oil and grease.

**Highlights:**

- Granular, biofilm and planktonic assemblages developed from the same seed reservoir
- Three assemblages forming metacommunity differed in diversity and composition
- Multiple null models applied to quantify deterministic and stochastic mechanisms
- Dominant, active role of acetoclastic *Methanosaeta* confirmed in granules and biofilm
- Dynamic, non-core-microbiome taxa correlated with environmental variables

## 1. INTRODUCTION

Microbial consortia have been exploited for numerous biotechnological applications, such as for valorizing wastewaters, principally because mixed-species systems allow for easier management and provide more diverse utility than pure-culture set-ups (McCarty and Ledesma-Amaro, 2019). The anaerobic digestion (AD) process, in particular, relies on the concerted activity of multiple microbial trophic groups to sequentially drive the bioconversion of organic matter to methane and CO_2_ in various assemblage modes – granular, biofilm or planktonic – and bioreactor configurations (Abdelgadir et al., 2014; Davey and O’Toole, 2000). The composition of the underlying methanogenic consortia responds to environmental and process selection pressures (Nemergut et al., 2013). Microbial taxa variously interact in forming community assemblages that are structured as biofilm slimes, flocs, granules and other aggregates, or, as planktonic communities (Aqeel et al., 2019; Davey and O’Toole, 2000). Thus, AD assemblages harbouring multiple trophic-level interactions thus serve as suitable environments to evaluate microbial community dynamics and assembly.

Unveiling microbial assembly mechanisms offers unprecedented insights into engineered bioconversion systems, such as AD bioreactors (Ferguson et al., 2018). Factors such as inoculum composition (Han et al., 2016; Li et al., 2019; Singh et al., 2019a), substrate composition and loading (Braz et al., 2019; Chen et al., 2019; Goux et al., 2015), operational duration (Lucas et al., 2015; Vanwonterghem et al., 2014), and process temperature (Heidrich et al., 2018), were shown as strong drivers of microbial community assembly in various AD systems when operated at hydraulic retention times (HRTs) longer than 10 d. However, shorter HRTs (<3 d) are sought to economize the anaerobic wastewater treatment process. Under such conditions, stochastic community assembly may be promoted due to random changes from immigration, drift or dispersal of microbial taxa (Nemergut et al., 2013; Stegen et al., 2012). Alternatively, short HRTs may promote environmental filtering due to the selection of taxa with unique survival potential (Stegen et al., 2012; Zhou, 2017). Treatment of organics, including inhibitory compounds at low (sub-mesophilic) temperatures is also desirable in AD systems to improve the net energy yield (Petropoulos et al., 2019), but such conditions are regarded as challenging for optimal functioning of AD consortia. Thus, operational conditions, including HRTs, the presence of inhibitory compounds, and operating temperatures, are likely drivers of microbial community assembly in methanogenic consortia. A mechanistic understanding of the concerted effects of these challenging growth conditions on the microbial community dynamics and assembly of methanogenic consortia will aid in further widening the applications of AD for waste bioconversion and valorisation.

A growing body of literature has characterized assembly and succession of microbial communities in granules (Liébana et al., 2019), biofilms (Xu et al., 2019) and planktonic assemblages (Xu et al., 2020) obtained from engineered microcosms, including more recently from AD bioreactors (Trego et al., 2021). However, AD bioreactors may rely on distinct assemblage modes to perform complementary roles across discrete reactor compartments (McAteer et al., 2020; Paulo et al., 2020; Singh et al., 2020). Thus, a systematic inference of the selective environmental pressures on the development of distinct assemblages from a common, seed community reservoir in AD bioreactors is needed to comprehend their comparative roles and assembly mechanisms.

The objective of this study was to characterize three distinct microbial assemblages (granules, biofilm, and planktonic communities) that were sampled from replicated bioreactors inoculated with one community reservoir of anaerobic sludge. The assemblages were developed in novel, two-compartment bioreactors treating long-chain fatty acids (LCFA)-rich wastewater at low HRTs (72-12 h) at 20°C, which we previously evaluated and reported (Singh et al., 2020). The temporal variations in active microbiomes from each of the assemblages were studied to evaluate the effect of environmental variables on microbial community dynamics (diversity, composition, and core and dynamic taxa). We applied multiple null model approaches to study community assembly, and to quantify the relative contribution of assembly processes involved in structuring the three assemblages.

## 2. MATERIALS AND METHODS

### 2.1 Sample collection

Microbial samples were collected from three, identical anaerobic dynamic sludge chamber-fixed film (DSC-FF) bioreactors, which were operated in parallel to treat mixed-LCFA-rich synthetic dairy wastewater at 20°C for 150 d as described previously (Singh et al., 2020). Granules from the sludge-bed layer (the DSC), the biofilm (the FF, which was grown on pumice stones) and the planktonic community from effluent samples were collected from different operating phases at the applied HRTs (72, 42.5, 24, 18, 12 h) corresponding to days 8, 24, 58, 100 and 148. The anaerobic sludge used as the inoculum was also sampled. Pumice stones were aseptically collected from the FF section, and sonicated for 5 min at 20°C with 10 mL of phosphate buffer saline (pH 7.2) to dislodge microbial cells from the attached biofilm. The supernatant from biofilm, and the effluent samples (25 mL), were each centrifuged at 8000 rpm for 10 min at 4°C to harvest cells. The resultant pellets, along with the granules and the inoculum samples, were flash-frozen in liquid nitrogen immediately upon collection and stored at −80°C.

### 2.2 Nucleic acids extraction and 16S rRNA gene amplicon sequencing

The microbial samples were thawed on ice, and the nucleic acids (DNA and RNA) were extracted using the phenol chloroform method (Griffiths et al., 2000). DNA and RNA concentrations were determined using a Qubit fluorometer (Life Technologies), and DNA purity was evaluated using a Nanodrop (NanoDrop Technologies, Wilmington, USA) and gel electrophoresis (Singh, 2019). Next, DNase treatment was performed to remove DNA using Invitrogen Turbo-DNase kit (Thermo Fisher, USA) by following the recommended procedure, and the DNA-free samples consisting of RNA were converted to cDNA using M-MuLV Reverse Transcriptase kit (New England BioLabs, USA) according to the instructions provided by the supplier. PCR amplification of the V4 region of the 16S rRNA gene was performed on cDNA transcripts with the universal primers 515f and 806r (Caporaso et al., 2011), with the sequencing of the active microbiomes on the Illumina MiSeq platform. Biofilm and effluent samples obtained from 72 h HRT were not sequenced because extractions yielded inadequate quantities of cDNA. The 16S rRNA sequences used to support the findings of this study have been deposited in the NCBI Sequence Read Archive under bioproject accession PRJNA657615.

### 2.3 Computational sequence analyses and statistical tests

The sequence data was analyzed using Quantitative Insights Into Microbial Ecology (QIIME v1.9) pipeline. The paired-end reads were joined using a fastq-join method with a min overlap of 50 bp and a perc_max_diff of 15%, after which quality filtering was performed using the split_libraries_fastq.py script in QIIME. The sequences were clustered into operational taxonomic units (OTUs) using the open-reference OTU picking with the BLAST method, and taxonomy assignment was performed against the Silva v132. It should be noted that chimeric sequences were identified using ChimeraSlayer (Haas et al., 2011) and the final OTU table was generated from the nonchimeric sequences using the script make_otu_table.py in QIIME (Aronesty, 2013; Caporaso et al., 2010). Statistical analyses were performed in R using all of the OTUs.

Quantification and visualization of microbial community diversity - alpha diversity (richness, Shannon indices), phylogenetic alpha diversity (Net Relatedness Index (NRI), Nearest Taxon Index (NTI), beta diversity (Principal Coordinate Analysis (PCoA) ordination plots), and, microbial community composition (25 most abundant classes and genera, and core microbiome) were performed in R. Representative taxa explaining beta diversity were obtained using the subset analysis to indicate the dynamic taxa in the dataset.

The core microbiome, considered at a prevalence of 85% across all samples was determined using the library microbiome (Lahti et al., 2019; Shetty et al., 2017). Subset analysis was performed to obtain the minimum set of representative OTUs that explained the beta diversity equivalent to using all the OTUs. The function bvStep() from the library sinkr evaluated the 2000 most abundant OTUs using the BVSTEP routine to achieve the highest possible correlation between dissimilarities of fixed and multivariate features through Mantel test (Clarke and Ainsworth, 1993; Taylor, 2017). Different combinations of the representative subset features were correlated to the full dataset, and, then PERMANOVA values for the combinations were calculated using 999 permutations. Redundancy analysis (RDA) on the Hellinger-transformed sequence data with forward selection (based on 999 permutations, variables retained at p<0.05) was performed to select for environmental variables most strongly associated with variance of the observed communities, indicating the variables structuring microbial communities. Correlation analysis was performed through bivariate analysis to measure the strength of association between representative OTUs (obtained from subset analysis) and the selected environmental variables, using Kendall correlation. Adjusted p-values were determined to assign the significance of the association.

### 2.4 Null model quantification of microbial assembly mechanisms and ecological processes

Different null modeling approaches with different analytical formulations were performed to achieve a general consensus on community assembly trends, and to mask out any biases associated with the methods. First, ecological stochasticity in community assembly based on beta diversity was calculated using NST with various distance measures (incidence-based: Jaccard, Kulczynski and abundance-based: Ruzcika, Kulczynski) in R using the NST package with 50% as the boundary point between more deterministic (<50%) and more stochastic (>50%) assembly (Liang et al., 2019; Ning et al., 2019; Zhou et al., 2014).

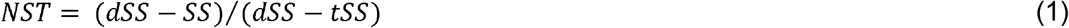

where SS is the observed selection strength, dSS and tSS are the theoretical extreme values of SS under completely deterministic and stochastic assembly, respectively. For the calculations, the taxa occurrence frequency and the sample richness were constrained as proportional (P) or fixed (F) in the combinations PF or PP, and 1000 randomizations were performed for each model. Statistical significance was computed based on permutational multivariate ANOVA (PANOVA). The choice of distance measures was based on the recommendations from a study where a simulated community with known stochasticity values was evaluated (Ning et al., 2019). Next, we used an alternative approach utilizing QPE analysis (Vass et al., 2020) to further explore the prevailing ecological niches, breaking down the community assembly mechanisms into the ecological processes to ‘homogenizing selection’, ‘variable selection’, ‘dispersal limitation’, ‘homogenizing dispersal’, and ‘undominated stochastic processes’. The ecological processes were quantified based on phylogenetic signal and abundance-based (Raup-Crick) beta-diversity (*β*_*RCBray*_) to discern amongst deterministic processes and stochastic processes following the framework of Stegen et al (2013) using beta mean nearest taxon distance (*βMNTD*) and beta nearest taxon index (*βNTI*). The f3*MNTD* value quantifies the phylogenetic distance between each OTU in one community and its closest relative in a second community.

## 3. RESULTS AND DISCUSSION

### 3.1 Microbial diversity patterns in granular, biofilm and planktonic assemblages

Microbial diversity (Fig. 1A,B) in the bioreactors varied with the applied HRT across the three assemblages (granules, biofilm and planktonic communities). Rarefied richness and Shannon entropies in the granules had not changed by the first sampling (day 8), but both were significantly (p<0.01) reduced when HRT was shortened from 72 to 12 h (Fig. 1A,B). This is in line with the previous studies using anaerobic microcosms, wherein niche specialization was linked to substrate-specific metabolic adaptations in the community (Braz et al., 2019). Meanwhile, in the biofilm, rarefied richness and Shannon entropies were initially lower than in the inoculum or the granules (Fig. 1A,B), but increased significantly (p<0.05) with the growing, *de novo* biofilm and with the changing HRT. Ultimately (by the time the HRT had been reduced to 12 h), the biofilm community was similarly diverse as, but more even (p<0.01) than in the granules or the planktonic microbiomes.

**Fig. 1.**
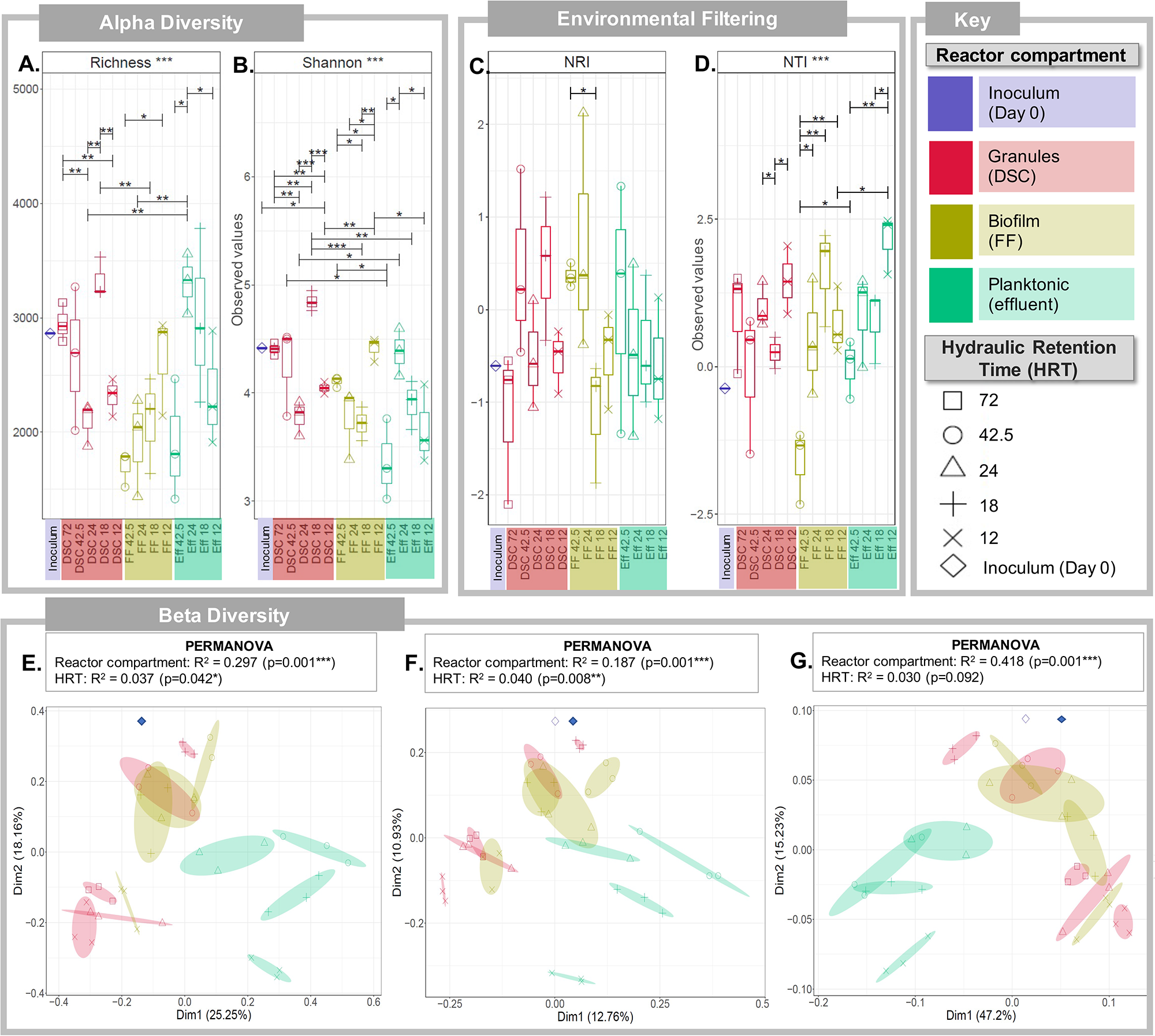
Diversity in the microbiomes of the inoculum and granular (from DSC), biofilm (from FF) and planktonic (from effluent) assemblages sampled at the different HRTs (72, 42.5, 24, 18 and 12 h). Alpha diversity box plot of the (A) rarefied richness and (B) Shannon entropy. Environmental filtering box plot of the (C) Net Relatedness Index (NRI) and (D) Nearest Taxa Index (NTI). Beta diversity: (E) PCoA plot and (F) beta dispersion calculated using Bray-Curtis distances; (G) PCoA plot and (H) beta dispersion calculated using unweighted UniFrac distances; and (I) PCoA plot calculated using weighted UniFrac distances. Assemblage groups are differentiated by colour, and the ellipses are drawn for each sample group at standard error of ordination points, where arrows mark the direction of change in the community structure between HRTs within assemblage groups that were found to be significant by beta dispersion. PERMANOVA explains significant variability in microbial community structure from different bioreactor compartments and at different HRTs. Lines for panels A, B, C and D connect two sample groups at statistically significant levels indicated by asterisks as * (p <0.05), **(p<0.01) or ***(p<0.001).

Phylogenetic alpha diversity analysis further differentiated the three assemblages, since NTI determined significant increase in phylogenetic clustering in the biofilm and planktonic assemblages when the HRT was reduced from 72 to 12 h (NTI >0, p<0.01) and in granular assemblages when the HRT was reduced from 18 to 12 h (NTI >0, p<0.01) (Fig. 1D). Environmental filtering (NTI >0) dominated the three assemblages at the different HRTs; and eventually at 12 h HRT, NTI followed the trend: effluent>granules>biofilm. Thus, NTI values support that deterministic processes (environmental variables) contributed majorly to structuring of the microbial community in the three assemblages during their temporal development. It is noteworthy to mention that NTI is preferred when assessing the presence of significant phylogenetic signal across short phylogenetic distances (Wang et al., 2013), and thus, was considered more useful than NRI due to the lack of substantial trait data in this dataset.

Assemblages when differentiated based on the beta diversity in PCoA plots (Fig. 1E,F,G) revealed an overlap among the granular and biofilm microbiomes at all the HRTs, whereas the planktonic microbiomes clustered distinctly apart. While a majority of the variations between categories was explained by the assemblages (18.7-41.8%) (p=0.001); the HRT further explained 3-4% of the variations between categories (Fig. 1E,F,G).

### 3.2 Core and dynamic taxa in the granular, biofilm and planktonic assemblages

Ten classes represented 90% of the inoculum community as well as the three assemblages (Fig. 2A), among which 24 genera were highly (>75%) represented (Fig. 2B). These 24 genera were also present in the core microbiome (Fig. 3) of the three assemblages, including the ubiquitous prevalence of *Lactococcus* and uncultured *Syntrophaceae* (prevalence >90% at reads >1000). Additionally, *Methanosaeta* and uncultured bacterium *Cloacimonadaceae* were prevalent in the granules and biofilms (Fig. 3), whereas *Pseudomonas* and *Acinetobacter* were more prevalent in the planktonic community (Fig. 3) – demonstrating that differentiation in the core microbiomes was assemblage-specific.

**Fig. 2.**
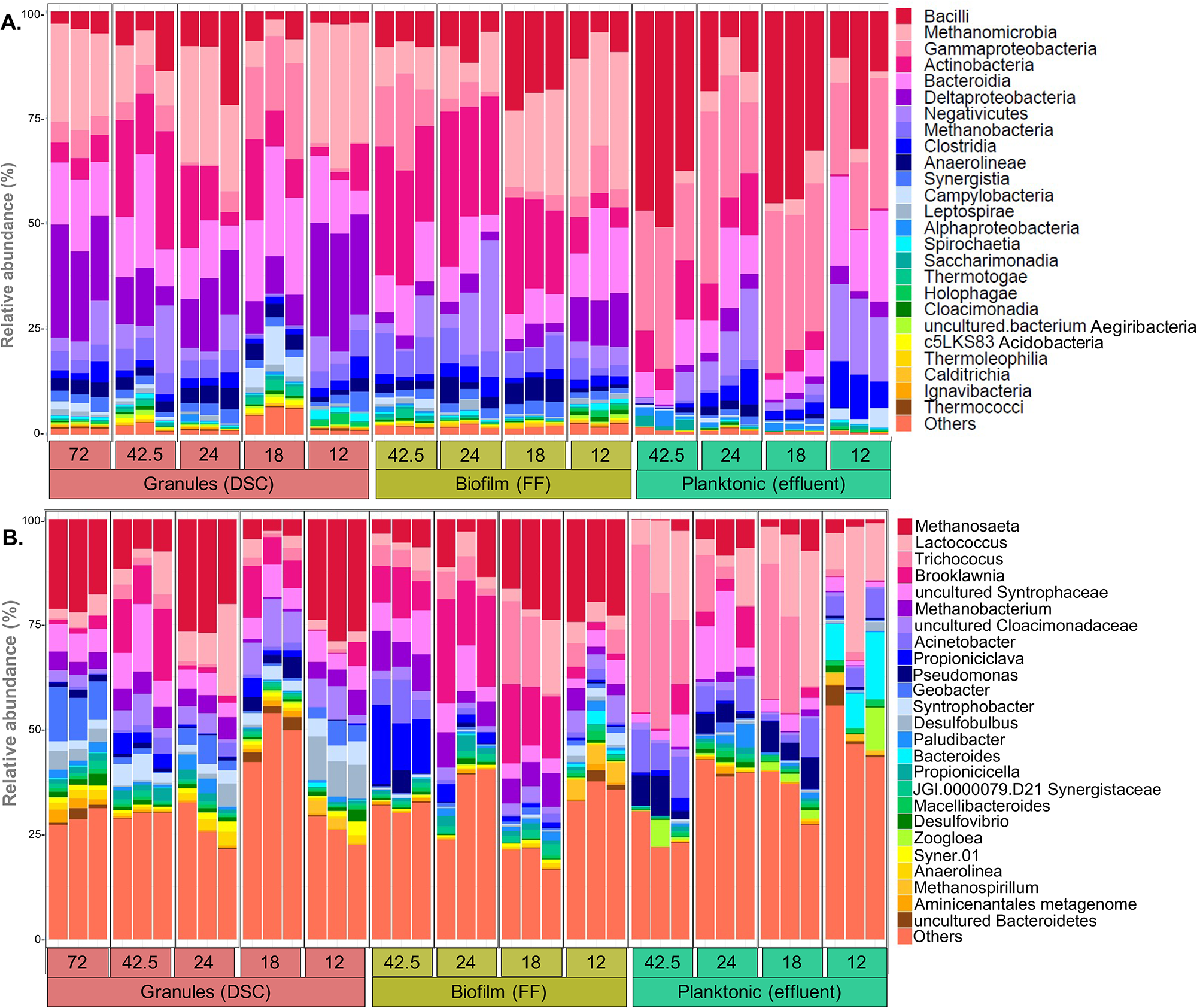
Barplots of the relative abundance of (A) the 25 most abundant classes, and (B) taxa identified to genus level, found in in the inoculum and the granular (obtained from the DSC), biofilm (obtained from FF), and planktonic (obtained from effluent) assemblages of the triplicate bioreactors R1, R2 and R3, at the HRTs of 72, 42.5, 24, 18 and 12 h. ‘Others’ are the taxa not included in the 25 most abundant.

**Fig. 3.**
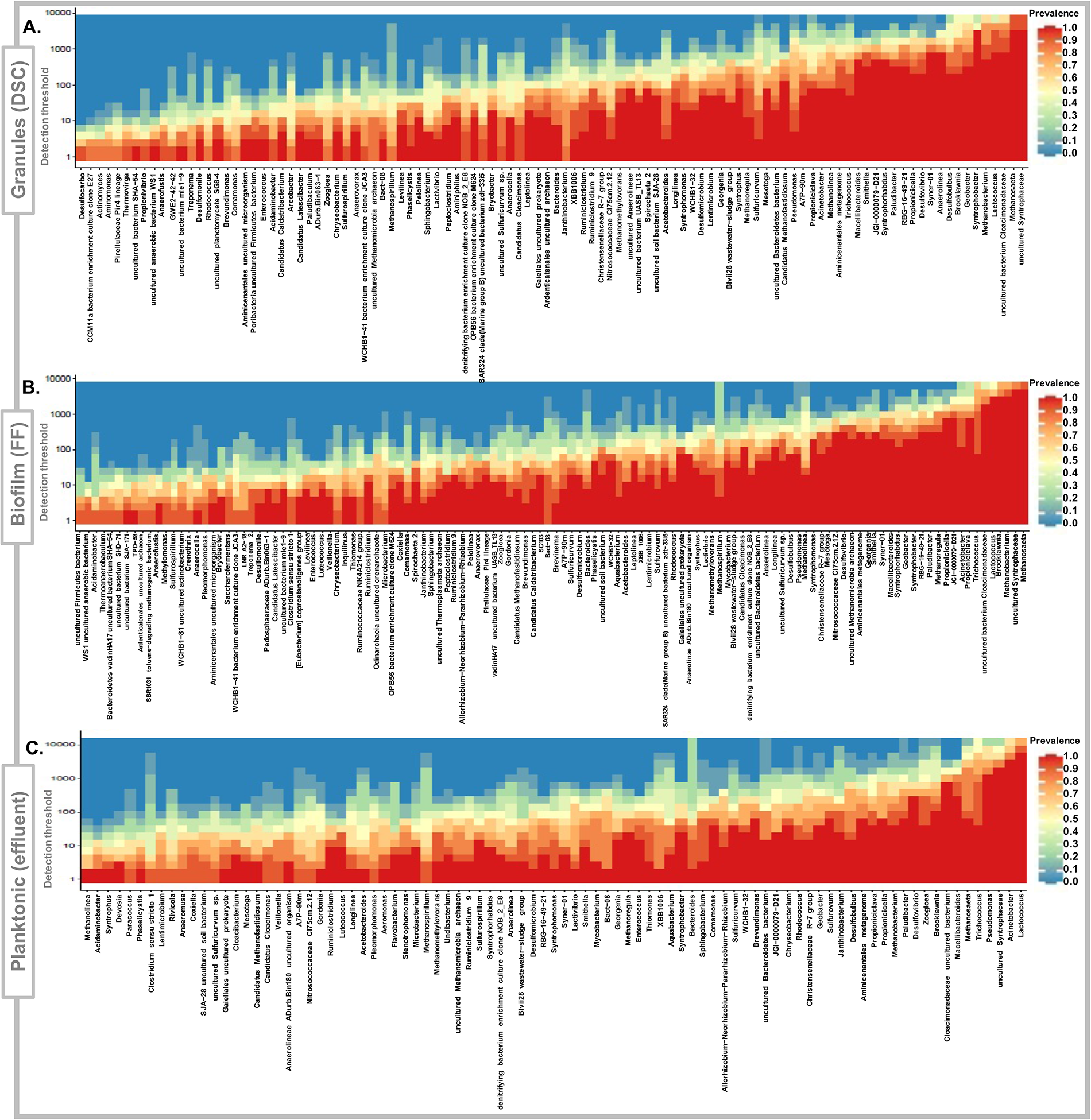
Highly prevalent taxa in the core microbiomes of (A) granular (from DSC), (B) biofilm (from FF), and (C) planktonic (from effluent) assemblages. Detection thresholds up to 10,000 reads are shown.

Subset analysis was performed to identify the minimum set of dynamic non-core taxa, namely the representative taxa that statistically explain the observed variances in the community (Table 1). Only five, six and nine taxa represented the variabilities in the beta-diversity of granular, biofilm and planktonic assemblages, respectively (Table 1). Thismeans a small fraction of the sequenced dataset explained 44.8-55.3%, 70-75.8% and 31.3-57.9% of the variation in, respectively, the granules, biofilms and planktonic microbial community dynamics at the decreasing HRTs in this study. Many of the core taxa (Fig. 3), such as *Methanosaeta*, *Methanobacterium*, uncultured *Syntrophaceae*, *Syntrophus*, *Syntrophobacter* and *Desulfobulbus,* have been previously found to be abundant during LCFA methanization at low ambient temperatures of 10-20°C (Grabowski et al., 2005; Singh et al., 2019a, 2019b). It is likely that these taxa played an active role at different trophic levels during the successive carbon flow during anaerobic LCFA methanization.

### 3.3 Quantifying mechanisms structuring the spatial development of assemblages

#### 3.3.1 Assembly mechanisms: Stochastic or deterministic?

We used a suite of null modelling approaches and quantified NST, *βNTI*, *βMNTD* and *β*_*RCBray*_ to determine the processes structuring microbial community assembly. The NST indices revealed significant differences in ecological stochasticity amongst the assemblages (p<0.001, PANOVA), irrespective of an evaluation through incidence-based or abundance-based distance matrices, or the constraints imposed on the taxa occurrence frequency and sample richness (Fig. 4A). NST indices were highest for the biofilm and lowest for the effluent (stochasticity trend: biofilm>granules>effluent) (Fig. 4A). QPE analysis revealed that the deterministic processes (55-61%) were higher than the stochastic processes (39-45%) in the assemblages. Trends in total stochasticity (‘dispersal limitation’, ‘homogenizing dispersal’, as well as the ‘undominated stochastic processes’) (Fig. 4B) found using QPE were similar to those found by the NST approach (Fig. 4), i.e, stochasticity followed the trend: biofilm>granules>effluent; and vice-versa for the deterministic causes. These results delineate that granular, biofilm and effluent assemblages have unique signatures, and stochastic and deterministic processes had combined roles in spatially structuring the community assembly in the distinct compartments of DSC-FF bioreactors.

**Fig. 4.**
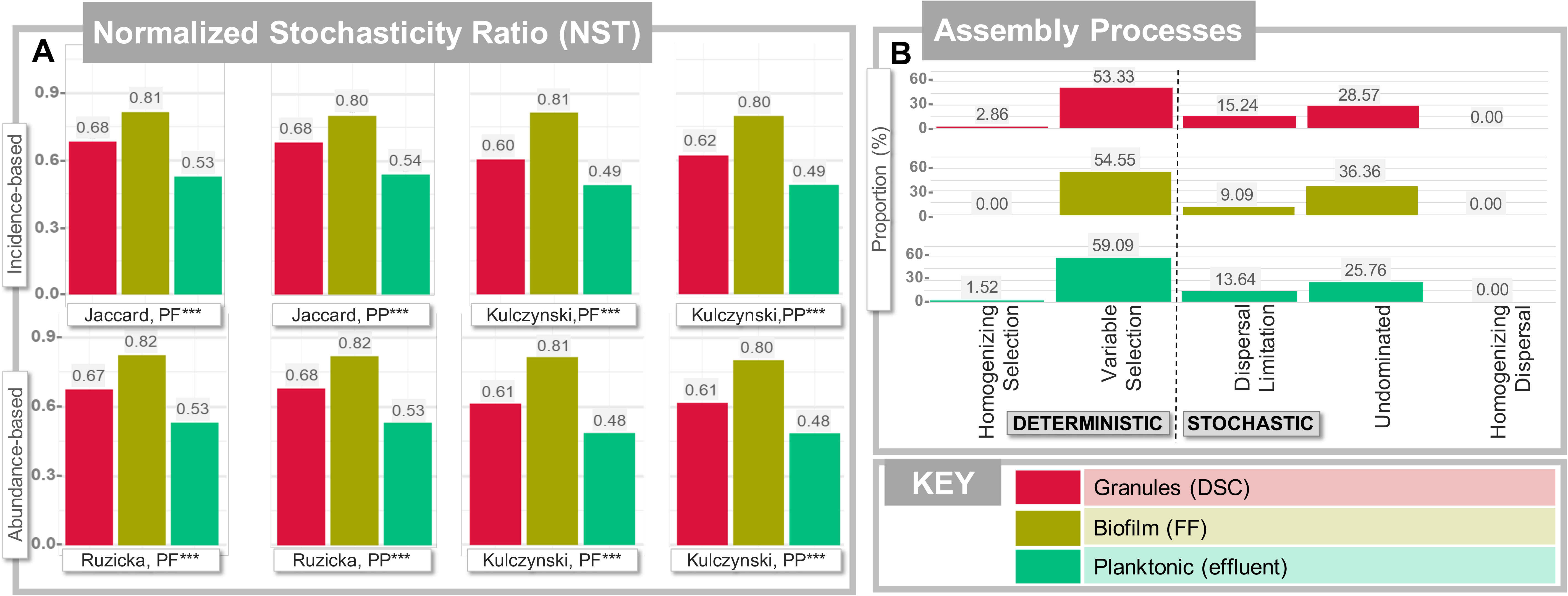
Estimates of microbial community assembly mechanisms structuring the spatial succession of granular (from DSC), biofilm (from FF) and planktonic (from effluent) assemblages grown from a single microbial community reservoir (inoculum). Spatial variation in (A) normalized stochasticity ratio (NST) indices, calculated as incidence-metrics (Jaccard and Kulczynski distance metrics) and as abundance-metrices (Ruzicka and Kulczynski distance metrices), with PF and PP null models among the granular, biofilm and planktonic assemblages; (B) the proportion of assembly processes by species-sorting (variable or homogeneous selection), dispersal limitation, homogenizing dispersal or undominated stochastic processes among the granular, biofilm and planktonic assemblages. Statistically significant levels are indicated by asterisks as *(p <0.05), **(p<0.01) or ***(p<0.001).

Amongst the deterministic causes, variable selection accounted for the largest proportion of the quantitative process (53-59%) (Fig. 4B), suggesting multiple niches for the selection of species which relied on the prevailing variability in environmental gradients. Dispersal limitation was the highest in the granules (15%), suggesting that among the sludge retention mechanisms (granulation and biofilm formation) in the DSC-FF reactors, granulation constrained dispersal more than biofilm formation. Overall, biofilm had the highest proportion of undominated stochastic processes (36%), and also occupied the highest proportion of all stochastic processes (45%) amongst the assemblages.

We used a comprehensive set of bioinformatics tools and null modelling approaches to quantify the diversity and assembly patterns in the AD assemblages to avoid possible analytical biases. The NST indices showed that both stochastic and deterministic processes shaped the granular, biofilm and planktonic assemblages under ‘challenging growth conditions’ in the triplicate bioreactors (Fig. 4A). NST is a relatively new measure that estimates stochasticity based on beta diversity measures, employing both incidence-based and abundance-based representations (Ning et al., 2019). Some of the null models also consider phylogenetic clustering as a proxy for the environmental drivers of community assembly. Therefore, looking at the community data with different null models using incidence and abundance matrices and phylogenetic data, our results show consistent assembly patterns (Fig. 4). Moreover, even the observed significant values of phylogenetic alpha diversity measured through NTI at 12 h HRT suggested the dominance of deterministic processes following the trend: effluent>granules>biofilm (Fig. 1D), which was consistent with the trends obtained from the NST and QPE estimations (Fig. 4).

Although conventional evaluations of microbial community assembly provided differentiation between deterministic and stochastic processes (Braz et al., 2019; Goux et al., 2015; Lucas et al., 2015), and null modeling approaches have been applied fairly recently for engineered bioreactors (Liébana et al., 2019; Santillan et al., 2019; Xu et al., 2020; Yuan et al., 2019), we were able for the first time to comprehensively explore microbial community structure and assembly from high-rate, anaerobic LCFA-treating bioreactors using multiple null models.

#### 3.3.2 Environmental variables affecting microbial community assembly

The three assemblages harbouring distinct regional microbial pools were linked hydraulically inside each of the respective bioreactors, and formed a metacommunity in each of the three separate bioreactors. Under short HRTs, microbes with low growth rates will ordinarily be washed out from suspended systems (i.e. those systems without retention on surfaces or in aggregates), leaving highly proliferating microbes as dominant populations (Yuan et al., 2019). Such effects are expected to be even more pronounced at low temperatures since the AD microbiome is considered optimally mesophilic (Pommerville, 2014), yielding diminished growth rates at reduced temperatures (Bergamo et al., 2009; Pavlostathis and Giraldo-Gomez, 1991; Petropoulos et al., 2018). Moreover, substrate characteristics strongly affect the microbial community structure. For example, LCFAs have a surfactant effect (Daffonchio et al., 1995) that leads to biofilm thinning and disaggregation of granular sludge (Arnaiz et al., 2003; Cavaleiro et al., 2001; Nikolaeva et al., 2013) and inhibit microbial activity (Astals et al., 2014). Thus, microbes less adept at adhering to biofilm or granular assemblages may be selectively washed out mobilizing microbes hydraulically within the metacommunity, particularly under short HRTs.

In this study, the stochastic processes could play a role either by immigration of taxa (e.g., from the granules in the DSC to the FF biofilm), or dispersal (washout) of taxa (e.g., from granules and the FF biofilm), or undominated stochastic processes leading to random changes in community composition in the assemblages. Concurrently, deterministic processes could play a role due to inter-species interactions (e.g., syntrophic interactions), or environmental variables (e.g., LCFA concentrations, HRT). While the relative contribution of each process has remained controversial (Stegen et al., 2012), we found that deterministic and stochastic mechanisms had combined roles in the microbial community assembly, wherein, biofilm had the highest stochasticity among the assemblages (Fig. 4). The bioreactors were designed as retained-biomass systems in this study so as to minimize the washout of slow-growing species. Morever, the biofilm compartment was designed to compensate for washout from the preceding sludge chamber (DSC). Microbes were washed upward from the variably-performing DSC chamber (granules) to the FF chamber (biofilm), and onwards to the effluent. We hypothesize that within the metacommunity, fluctuating metabolite (substrate and intermediates) concentrations and microbial populations reached the biofilm from DSC, which resulted in a higher randomness in the biofilm than in the granules or effluent microbiomes.

As a variable selection accounted for the largest proportion (>50%) of the assembly processes, we considered it interesting to identify the environmental variables significantly linked to the variability in the observed communities. RDA with forward selection was employed (Vass et al., 2020), which showed that the stearate loading rate (p<0.001), palmitate loading rate (p<0.01), palmitate removal rate (p<0.05) and caproate concentrations (p<0.01) had a bearing on the variation in microbial community composition (Table 2). Specifically, this study could associate microbial community assembly with operational parameters (HRTs, and LCFA loading (stearate and palmitate)) (Table 2). These findings have transferrable implications to reactor operation strategies and could be employed to modulate microbial community assembly in anaerobic bioreactors operated at conditions similar to this study. For example, higher dispersal limitation was observed in granules than biofilms (Fig. 4), reflecting the utility of granulation in microbial retention under challenging operational conditions.

### 3.4 Environmental effects on microbial community dynamics

#### 3.4.1 Environmental variables affecting microbial community composition and dynamics

Given that the microbial community diversity, composition and assembly were differentiated in the assemblages and demonstrated predominant environmental filtering, we investigated the environmental variables that delineated the microbial community dynamics by applying correlation analysis. The concentrations of total and soluble COD, caproate (C_6_), palmitate (C_16_) and total LCFAs were strongly correlated (p <0.05) to the taxa abundance (Fig. 5). The strength of association between the representative taxa and environmental variables was measured using Kendall correlation. Palmitate removal rate positively correlated to the representative taxa, *Geobacter*, *uncultured bacterium Syner-01*, *uncultured bacterium SJA-29* and *Syntrophobacterales bacterium Delta 03*. Of these, *uncultured bacterium Syner-01*, and *uncultured bacterium SJA-29* were negatively correlated to the concentrations of even-chained VFAs, specifically acetate (C_2_) or butyrate (C_4_). Whilst the *uncultured bacterium WCHB1-32* positively correlated to tCOD, sCOD and caproate (C_6_) concentrations, the *uncultured bacterium WCHB1-41* negatively correlated with the loading rates of palmitate (C_16_) and stearate (C_18_), thereby delineating the roles of different fatty acids in structuring the dynamic non-core taxa in the assemblages.

**Fig. 5.**
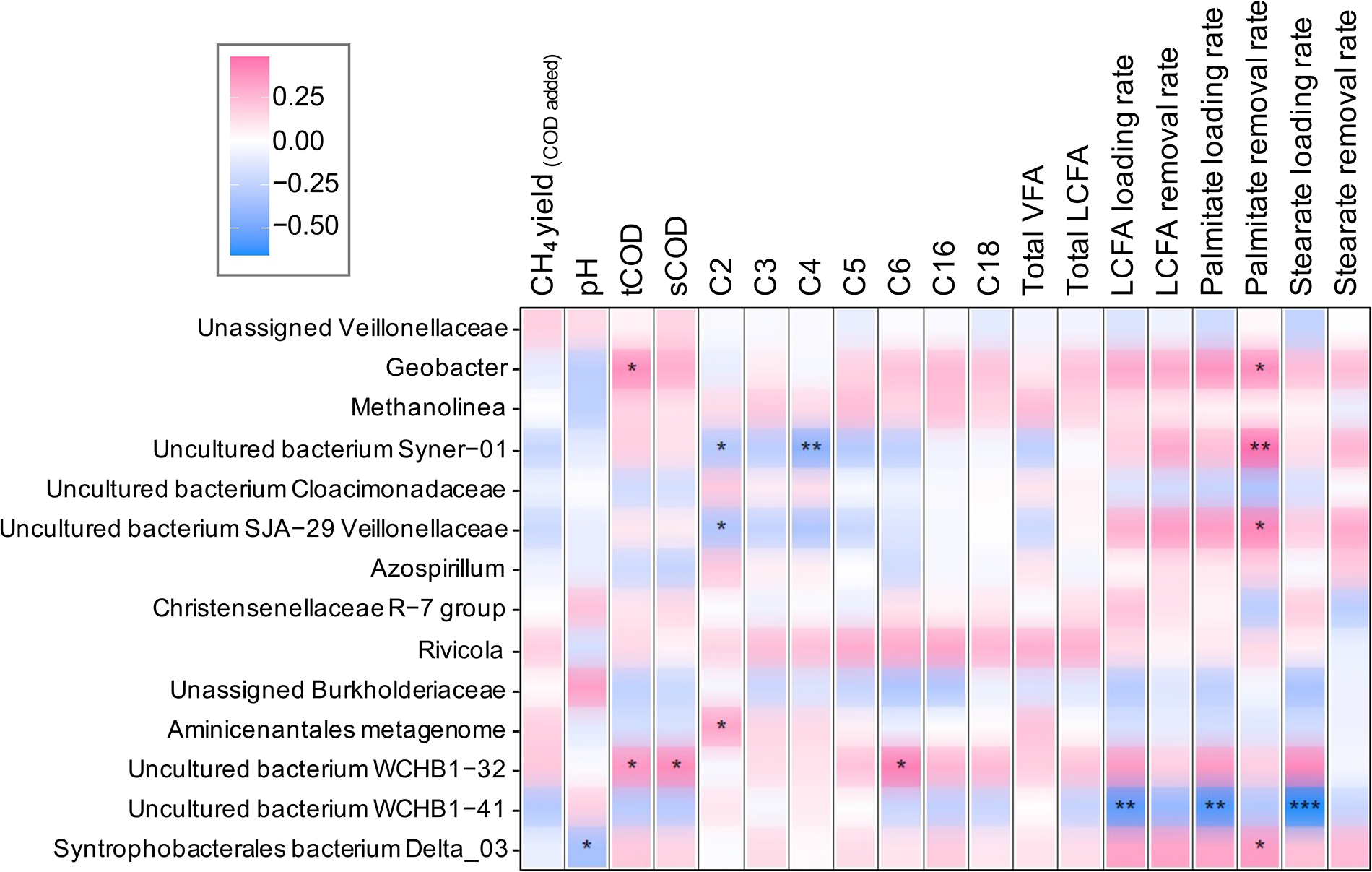
The correlation heatmap of the representative taxa from granular (from DSC), biofilm (from FF) and planktonic (from effluent) assemblages with the environmental variables (metabolite concentrations and process performance parameters) prevailing in the different bioreactor compartments (DSC, FF, effluent). Kendall correlations between the representative taxa and the environmental variables were calculated. Significance levels are indicated by asterisks as *(p <0.05), **(p<0.01) or ***(p<0.001).

#### 3.4.2 Anaerobic LCFA degradation by assemblages: methanogenesis and syntrophic β-oxidation

HRTs and LCFA loading rates influenced the representative taxa driving community dynamics in the granular, biofilm and planktonic assemblages (Fig. 5, Table 1). With decreasing HRTs and LCFA loading, equivalently increasing active populations of fermenters, β-oxidizers, and hydrogenotrophic and acetoclastic methanogens were required for degradation of the LCFAs at 20°C. Methanogenesis is a conserved function mediated through methylotrophic, acetoclastic or hydrogenotrophic pathways. Among the archaeal taxa found in the granular sludge microbiome in this study, *Methanosaeta* was the only acetoclastic genus, and had the highest relative abundances as well as high prevalence in core microbiomes (Fig. 2,3). Earlier, *Methanosaeta* was found in high relative abundances in LCFA-degrading anaerobic consortia in both batch assays and continuous high-rate reactors at low temperatures (Lv et al., 2015; Paulo et al., 2020; Singh et al., 2019a, 2019b). In the current study, by analyzing 16S rRNA (cDNA) we could demonstrate the persistence and active role of *Methanosaeta* in granules and biofilm in the acetoclastic methanogenesis at 20°C even at low HRTs and high LCFA loading.

β-oxidation of LCFAs is a narrowly conserved function, wherein only few known species belonging to the genera *Syntrophus* (family *Syntrophaceae,* class *Deltaproteobacteria*)*; Syntrophomonas, and Thermosyntropha* (family *Syntrophomonadaceae,* class *Clostridia*), and uncultured taxa (family *Clostridiaceae*, class *Clostridia*) (Baserba et al., 2012; Sousa et al., 2009). Thus, an examination of taxa belonging to the families *Syntrophomonadaceae* (class *Clostridia*) and *Syntrophaceae* (class *Deltaproteobacteria*) in this study was considered of interest to find taxa responsible for LCFA degradation. In our study, the relative abundance of *Syntrophomonas* was low (<0.5%), whereas taxa assigned to the family *Syntrophaceae* were highly abundant, consisting of *Syntrophus* and an uncultured *Syntrophaceae* taxon (Fig. 2). In previous studies, *Syntrophomonadaceae-*related taxa have been reported frequently from mesophilic or thermophilic anaerobic bioreactors treating fats, oils and grease (FOG)-rich wastes at relative abundances, that is, at 0.2-25% in sludges treating lipid-rich wastes (Hansen et al., 1999; Menes and Travers, 2006; Ziels et al., 2017). Concurrently, *Syntrophus-*affiliated taxa have been reported from diverse environments, including not only the mesophilic and thermophilic digesters treating palm-oil mill effluent (POME) and LCFAs (Hatamoto et al., 2007; Yoochatchaval et al., 2011), and mesophilic reactors treating cafeteria wastewater (Fujihira et al., 2018); but also the psychrophilic digesters treating food waste (Choudhary et al., 2020) and LCFA-rich wastewater (Singh et al., 2019a), low-temperature reactors treating LCFA-rich wastewater (Singh et al., 2019b), and psychrophilic digesters degrading stearate and alkanes (Grabowski et al., 2005). Meanwhile, highly abundant populations of both *Syntrophus* and *Syntrophomonas* have also been reported, e.g., from batch incubations digesting lipid-rich scum at 30°C wherein the relative abundances increased for *Syntrophus* (3-fold) as well as *Syntrophomonas* (1.1-fold) (Fujihira et al., 2018). Thus, linking the prevalence of the β-oxidizing genera to the operational conditions has been confounding based on substrate (FOG, LCFA) and operational conditions (temperature, HRTs).

In our study, uncultured *Syntrophaceae* was the only putative β-oxidizer present at high relative abundances (Fig. 2) as well as in the core microbiomes (Fig. 3). Hence, an active role for *Syntrophaceae*-affiliated taxa (*Syntrophus* and uncultured) is implied in treating LCFA-rich wastewaters at low HRTs at 20°C; supported by their psychrotolerant growth in granules as well as in biofilms (Fig. 2,3). Syntrophs are fastidious and despite playing a crucial role in global carbon cycling, their characterization has remained relatively obscure (Narihiro and Kamagata, 2017). Although taxonomic resolution of uncultured syntrophic bacteria below family level may still be challenging despite the use of high-end molecular methods (Hatamoto et al., 2007), we show that the use of new bioinformatics approaches offers alternative ways to link process performance with population dynamics in microbial communities. This may provide useful insights when low microbial growth rates, or an abundance of uncultured taxa, confound analyses using conventional molecular approaches. The large proportion of unassigned and uncultured taxa in our study suggests that novel uncultured microbes may have a role in methanization of LCFAs in the granular and biofilm assemblages at low ambient temperatures. Future advancements should leverage genome-centric tools to connect the abundance of β-oxidizing bacteria to LCFA metabolism in response to diverse external stimuli, while simultaneously comprehending the functional expressions of *Syntrophus* and *Syntrophomonas* under those stimulus (James et al., 2019; Sieber et al., 2015; Treu et al., 2016).

## 4. Conclusions

Multiple null modelling approaches systematically confirmed that combined deterministic and stochastic mechanisms influenced the microbial community assembly in high-rate bioreactors treating LCFA-rich wastewater at 20ºC. Variable selection (deterministic) accounted for the largest proportion (>50%) of the assembly processes, while undominated processes (26-36%) constituted the most important stochastic process. Abundant *Methanosaeta* and *Syntrophaceae* (*Syntrophus* and uncultured taxa) were prevalent in the core microbiomes, suggesting these taxa were critical in syntrophic LCFA-degradation. This study expands our understanding of the microbial community dynamics and assembly in complex metacommunities comprising granular, biofilm and planktonic assemblages mediating dairy waste bioconversion in innovative DSC-FF bioreactors.

## Data availability

The sequencing data from this study are available through the NCBI database under the project accession number accession PRJNA657615.

## CRediT authorship contribution statement

**SS**: Conceptualization, Investigation, Data curation, Formal analysis, Writing - original draft, Writing - review & editing. **JMRK**: Writing - review & editing. **PNLL**: Funding acquisition, Writing - review & editing. **MK**: Writing - original draft, Writing - review & editing. **JR**: Funding acquisition, Writing - review & editing. **VOF**: Conceptualization, Supervision, Writing - review & editing. **UZI**: Formal analysis, Supervision, Writing - review & editing. **GC**: Funding acquisition, Conceptualization, Supervision, Writing - original draft, Writing - review & editing.

## Declaration of Competing Interest

The authors declare they have no conflicts of interest.

## Acknowledgements

This project has received funding from the European Union’s Horizon 2020 research and innovation programme under the Marie Sklodowska-Curie European Joint Doctorate (EJD) in Advanced Biological Waste-To-Energy Technologies (ABWET), under grant agreement No 643071. VOF was supported by the Enterprise Ireland Technology Centres Programme (TC/2014/0016) and Science Foundation Ireland (14/IA/2371 and 16/RC/3889). UZI was funded by a NERC Independent Research Fellowship (NE/L011956/1) and EPSRC GCRF Grant (EP/P029329/1). GC, and DNA sequencing, was supported by a Science Foundation Ireland Career Development Award (17/CDA/4658).

## Supplementary data

Supplementary data to this article has been attached.

## Table Captions

Table 1. Correlation of representative taxa (obtained from subset analysis) to the full operational taxonomic unit (OTU) table.

Table 2. Redundancy analysis (RDA) with forward selection using ADONIS to select the environmental variables most strongly associated with the variance of the observed communities.

## Supplementary Figure Captions

Fig. S1. Schematic representation of the granular, biofilm and planktonic assemblages in the dynamic sludge chamber fixed-film (DSC-FF) bioreactors.

## Supplementary Table Captions

Table S1. Summary of operational conditions during synthetic dairy wastewater (SDW) treatment in the dynamic sludge chamber fixed-film (DSC-FF) bioreactors.

## Notes

### Competing Interest Statement

The authors have declared no competing interest.

