## Supplementary Material for "ASSEMBLY AND DYNAMICS OF MICROBIAL COMMUNITIES IN GRANULAR, FIXED-BIOFILM AND PLANKTONIC METHANOGENIC MICROBIOMES VALORIZING LONG CHAIN FATTY ACID (LCFA)-RICH WASTEWATER"

**List of Supplemental Material**

**Supplementary Text**. Statistical analysis.

**Figure S1**. Schematic representation of the granular, biofilm and planktonic assemblages in the dynamic sludge chamber fixed-film (DSC-FF) bioreactors.

**Table S1.** Table S1. Summary of operational conditions during synthetic dairy wastewater (SDW) treatment in the dynamic sludge chamber fixed-film (DSC-FF) bioreactors.

### Supplementary Text

**Statistical analysis**

***Alpha diversity metrices.*** Statistical analyses were performed in R using the tables and data generated as above as well as the meta data associated with the study. The Vegan package was used for the analysis of alpha and beta diversity in microbial samples (Oksanen et al., 2016). For alpha diversity, the indices used were: (i) rarefied richness – to represent the estimated number of species in a rarefied sample (to minimum library size); (ii) Shannon entropy – to represent a measure of balance within a community. R's aov() function was used to calculate the pair-wise analysis of variance (ANOVA) p-values which were then drawn on top of alpha diversity figures.

***Phylogenetic alpha diversity analysis: Environmental filtering and overdispersion.*** Further characterization of the community diversity was performed by the calculation of phylogenetic dispersion using: (i) nearest taxa index (NTI), and, (ii) net relatedness index (NRI), to evaluate the predominant microbial community assembly mechanism. The mean phylogenetic diversity (MPD) and functions mpd() and ses.mpd() were computed for NRI estimation, whereas the mean nearest taxon distance (MNTD) and functions mntd() and ses.mntd() were computed for NTI estimation by randomly generating null distributions with 1000 iterations in the library picante (Kembel et al., 2010), based on the presence or absence of taxa. NTI and NRI represent the negatives of the output from ses.mntd() and ses.mpd(), respectively. While NRI is a measure of mean pairwise phylogenetic distance of taxa in a sample, relative to phylogeny of an appropriate species pool and quantifies overall clustering of taxa on a tree; NTI is a measure of mean pairwise phylogenetic distance at local level and quantifies tip-level divergences (putting more emphasis on terminal clades and is akin to “local” clustering) in phylogeny. Hence, NTI is preferred over NRI when assessing presence of significant phylogenetic signal across short phylogenetic distances (Wang et al., 2013), or in cases where phylogenetic signal cannot be measured due to lack of substantial trait data. A significantly positive NTI value indicated that co-occurring species were more closely clustered than expected by chance, and vice-versa for the negative values. For a single community, NTI values > +2 suggest environmental filtering, and values < -2 indicate competitive exclusion among species as the driver of community structure (Stegen et al., 2012). A mean NTI taken across all communities that is significantly different from zero indicates clustering (NTI>0) or overdispersion (NTI<0) on average (Kembel, 2009). Based on the hypothesis that closely related taxa were more ecologically or functionally similar (phylogenetic niche conservatism), the obtained NTI measure can be used to infer ecologically similar (phylogenetic clustering) or ecologically dissimilar (phylogenetic overdispersion) taxa within a given community (Webb et al., 2002), and, was used to assess the influence of environment on microbial community structure in assemblages by quantifying the local clustering in the phylogenetic tree.

***Beta diversity and beta dispersion.*** For beta diversity, the dissimilarity in species community composition between pairwise comparisons of bacterial communities were represented in Principal Coordinate Analysis (PCoA) ordination plots by calculating three different distance metrices (Vegan ’s cmdscale() function): (i) Bray Curtis, considers the species abundance count; (ii) Unweighted Unifrac considers the phylogenetic distance between the branch lengths of operational taxonomic units (OTUs) observed in different samples, calculated using the phyloseq package (McMurdie and Holmes, 2013), and, (iii) Weighted Unifrac, considers the species abundance count with the phylogenetic distance. The samples were grouped for the conditions, mean ordination values and spread of points (standard errors of the (weighted) averages as ellipse) using Vegan’s CovEllipse () function.

The statistical scripts and workflows for all above can be found at:

<http://userweb.eng.gla.ac.uk/umer.ijaz#bioinformatics>.

### Supplementary Figure

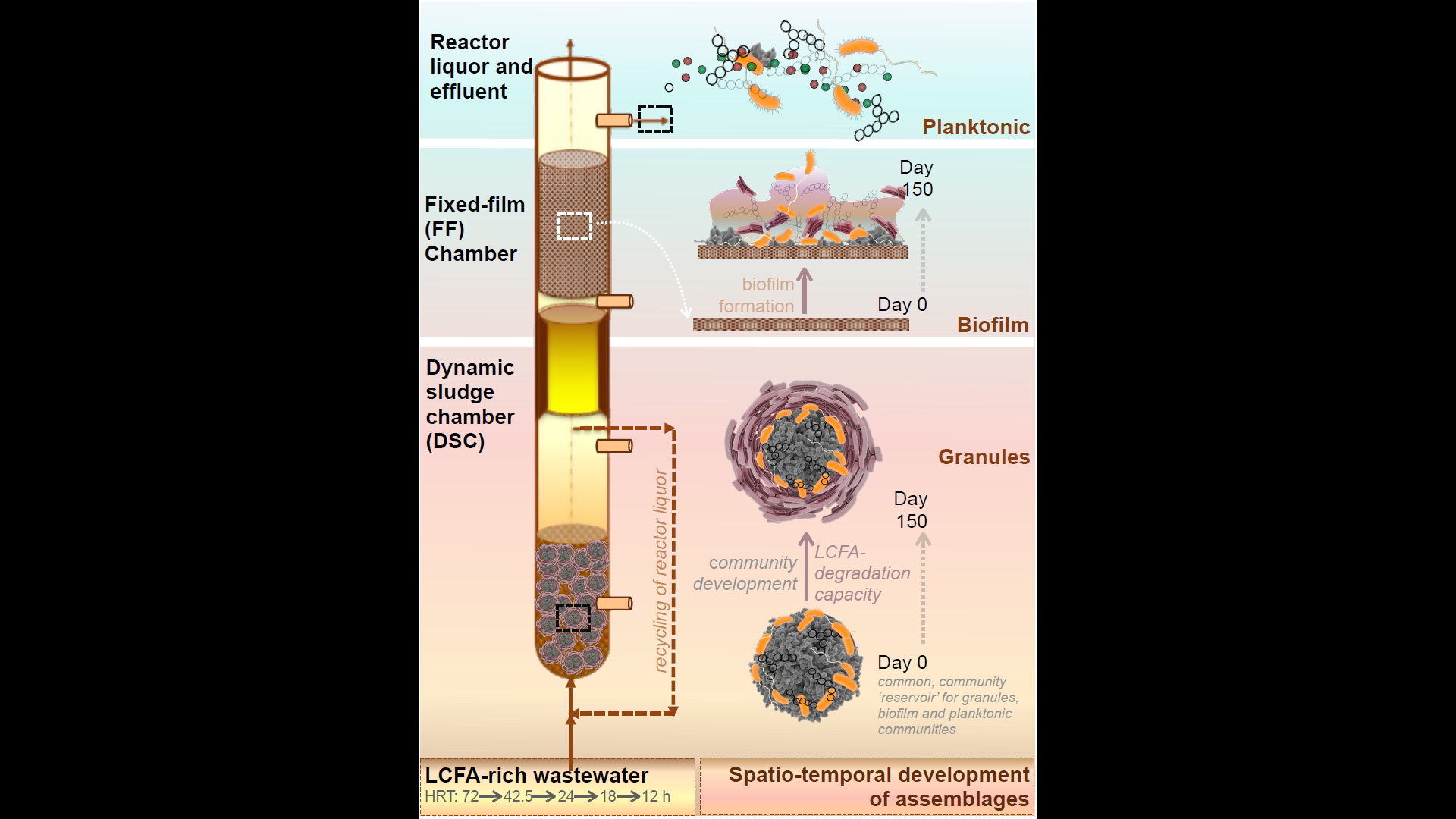

**Figure S1.** Schematic representation of the granular, biofilm and planktonic assemblages in the dynamic sludge chamber fixed-film (DSC-FF) bioreactors.

### Supplementary Table

Table S1. Summary of operational conditions during synthetic dairy wastewater (SDW) treatment in the dynamic sludge chamber fixed-film (DSC-FF) bioreactors.

| HRT (h) | **72** | **42.5** | **24** | **18** | **12** |
| --- | --- | --- | --- | --- | --- |
| pH | 7±0.1 | 7±0.1 | 7±0.1 | 7±0.1 | 7±0.1 |
| OLR (gCOD/L.d) | 0.66 | 1.13 | 2 | 2.67 | 4 |
| LCFA loading rate (gCOD-LCFA/L.d) | 0.22 | 0.37 | 0.67 | 0.89 | 1.33 |
| LCFA loading rate (mgCOD-LCFA/g-VS.d) | 34 | 68 | 120 | 180 | 240 |

HRT – Hydraulic retention time

OLR – Organic loading rate

LCFA – Long chain fatty acid
