## Supplementary material for "ASSEMBLY AND DYNAMICS OF MICROBIAL COMMUNITIES IN GRANULAR, FIXED-BIOFILM AND PLANKTONIC METHANOGENIC MICROBIOMES VALORIZING LONG CHAIN FATTY ACID (LCFA)-RICH WASTEWATER": Tables 1 and 2

Table 1. Correlation of representative taxa (obtained from subset analysis) to the full operational taxonomic unit (OTU) table.

| **No** | **Granular assemblage (DSC)** | **Correlation with full OTU table** | **PERMANOVA** |
| --- | --- | --- | --- |
| 1 | *Rivicola + uncultured bacterium Cloacimonadaceae + uncultured Burkholderiaceae + uncultured bacterium WCHB1-41 + uncultured Syntrophobacterales bacterium Delta_03* | 0.951 | R2 = 0.553;  p = 0.004 |
| 2 | *Rivicola + uncultured bacterium Cloacimonadaceae + uncultured Burkholderiaceae + uncultured Syntrophobacterales bacterium Delta_03* | 0.943 | R2=0.551;  p=0.003 |
| 3 | *Rivicola + uncultured bacterium Cloacimonadaceae + uncultured Syntrophobacterales bacterium Delta_03* | 0.9301 | R2=0.551; p=0.004 |
| 4 | *Rivicola + uncultured Syntrophobacterales bacterium Delta_03* | 0.914 | R2=0.448; p=0.067 |
|  | **Biofilm assemblage (FF)** | **Correlation with full OTU table** | **PERMANOVA** |
| 1 | *Methanolinea + uncultured bacterium SJA-29 + Geobacter + Christensenellaceae R-7 group + uncultured Veillonellaceae* | 0.833 | R^2^=0.75639; p=0.003 |
| 2 | *Methanolinea + Methanoregula + uncultured bacterium SJA-29 + Geobacter + Christensenellaceae R-7 group + uncultured Veillonellaceae* | 0.828 | R^2^=0.701; p=0.008 |
| 3 | *Methanolinea + Geobacter + Christensenellaceae R-7 group + uncultured Veillonellaceae* | 0.82 | R^2^=0.758; p=0.002 |
| 4 | *Methanolinea + Geobacter + Christensenellaceae R-7 group* | 0.809 | R^2^=0.758; p=0.001 |
| 5 | *Methanolinea + Christensenellaceae R-7 group* | 0.799 | R^2^=0.758; p=0.002 |
| **No** | **Planktonic assemblage (effluent)** | **Correlation with full OTU table** | **PERMANOVA** |
| 1 | *Rivicola* + *uncultured bacterium Cloacimonadaceae + uncultured Burkholderiaceae + Methanolinea + Aminicenantales metagenome + uncultured bacterium WCHB1-32 + Syner-01 uncultured bacterium + uncultured Syntrophobacterales bacterium Delta_03 + Azospirillum* | 0.951 | R^2^=0.57803; p=0.001 |
| 2 | *Rivicola + uncultured bacterium Cloacimonadaceae + uncultured Burkholderiaceae + Methanolinea + Aminicenantales metagenome + uncultured bacterium WCHB1-32 + Syner-01 uncultured bacterium + uncultured Syntrophobacterales bacterium Delta_03* | 0.938 | R^2^=0.57854; p=0.001 |
| 3 | *Rivicola + uncultured bacterium Cloacimonadaceae + uncultured Burkholderiaceae + Methanolinea + Aminicenantales metagenome + uncultured bacterium WCHB1-32 + Syner-01 uncultured bacterium* | 0.927 | R^2^=0.57928; p=0.001 |
| 4 | *Rivicola + uncultured bacterium Cloacimonadaceae + uncultured Burkholderiaceae + Methanolinea + Aminicenantales metagenome + uncultured bacterium WCHB1-32* | 0.903 | R^2^=0.485; p=0.021 |
| 5 | *Rivicola + uncultured bacterium Cloacimonadaceae + uncultured Burkholderiaceae + Aminicenantales metagenome + uncultured bacterium WCHB1-32 + Syner-01 uncultured bacterium* | 0.889 | R^2^=0.584; p=0.001 |
| 6 | *Rivicola + uncultured bacterium Cloacimonadaceae + uncultured Burkholderiaceae + Aminicenantales metagenome + uncultured bacterium WCHB1-32* | 0.867 | R^2^=0.494; p=0.017 |
| 7 | *Rivicola + uncultured bacterium Cloacimonadaceae + Aminicenantales metagenome + uncultured bacterium WCHB1-32 + Syner-01 uncultured bacterium* | 0.856 | R^2^=0.584; p=0.004 |
| 8 | *Rivicola + uncultured bacterium Cloacimonadaceae + Aminicenantales metagenome + uncultured bacterium WCHB1-32* | 0.827 | R^2^=0.479; p= 0.012 |
| 9 | *uncultured bacterium Cloacimonadaceae + Aminicenantales metagenome + uncultured bacterium WCHB1-32 + Syner-01 uncultured bacterium* | 0.807 | R^2^=0.521; p=0.029 |
| 10 | *uncultured bacterium Cloacimonadaceae + Aminicenantales metagenome + uncultured bacterium WCHB1-32* | 0.784 | R2=0.327; p= 0.297 |
| 11 | *uncultured bacterium Cloacimonadaceae + Aminicenantales metagenome + Syner-01 uncultured bacterium* | 0.762 | R^2^=0.525; p=0.031 |
| 12 | *uncultured bacterium Cloacimonadaceae + Aminicenantales metagenome* | 0.722 | R^2^=0.313; p=0.291 |
| 13 | *uncultured bacterium Cloacimonadaceae + Syner-01 uncultured bacterium* | 0.722 | R^2^=0.523; p=0.022 |

Table 2. Redundancy analysis (RDA) with forward selection using ADONIS to select the environmental variables most strongly associated with the variance of the observed communities.

| **Environmental variables** | **Df** | **SumsOfSqs** | **MeanSqs** | **F.Model** | **R^2^** | **Pr(>F)** | **Significance** |
| --- | --- | --- | --- | --- | --- | --- | --- |
| Stearate (C18:0) loading rate | 1 | 0.6644 | 0.66441 | 6.1819 | 0.14982 | 0.0001 | *** |
| Methane yield _(COD added)_ | 1 | 0.3466 | 0.34657 | 3.2246 | 0.07815 | 0.0039 | ** |
| Caproate (C6) concentrations | 1 | 0.3683 | 0.36828 | 3.4266 | 0.08305 | 0.004 | ** |
| Valerate (C5) concentrations | 1 | 0.2945 | 0.29448 | 2.74 | 0.06641 | 0.0141 | * |
| Palmitate (C16:0) loading rate | 1 | 0.3405 | 0.34051 | 3.1683 | 0.07679 | 0.0046 | ** |
| Palmitate (C16:0) removal rate | 1 | 0.2708 | 0.27084 | 2.52 | 0.06108 | 0.0201 | * |
| Residuals | 20 | 2.1495 | 0.10748 | 0.48471 |  |  |  |
| Total | 26 | 4.4346 | 1 |  |  |  |  |

Significance codes: 0 ‘***’ 0.001 ‘**’ 0.01 ‘*’ 0.05 ‘.’ 0.1 ‘ ’ 1
